# The Stress-Chip: A microfluidic platform for stress analysis in *Caenorhabditis elegans*

**DOI:** 10.1101/285163

**Authors:** Stephen A. Banse, Benjamin W. Blue, Kristin J. Robinson, Cody M. Jarrett, Patrick C. Phillips

**Author notes:** Current Address: Molecular Medicine and Mechanisms of Disease, University of Washington, Seattle, Washington, United States of America (Benjamin W. Blue). These authors contributed equally to this work (Stephen A. Banse, Benjamin W. Blue).

## Abstract

An organism’s ability to mount a physiological response to external stressors is fundamental to its interaction with the environment. Experimental exploration of these interactions benefits greatly from the ability to maintain tight control of the environment, even under conditions in which it would be normal for the subject to flee the stressor. ere we present a nematode research platform that pairs automated image acquisition and analysis with a custom microfluidic device. This platform enables tight environmental control in low-density, single-worm arenas, which preclude animal escape while still allowing a broad range of behavioral activities. The platform is easily scalable, with two 50 arena arrays per chip and an imaging capacity of 600 animals per scanning device. Validating the device using dietary, osmotic, and oxidative stress indicates that it should be of broad use as a research platform, including eventual adaptation for additional stressors, anthelmintic-drug screening, and toxicology studies.

## Introduction

The nematode *Caenorhabditis elegans* is a popular research model for studying a broad range of environmental effects, including osmotic, dietary, oxidative, hypoxia, heat, and heavy metal stress [1]. In most stress assays, resistance or susceptibility is measured either as the time until death post exposure or the percentage surviving in a population at set time points post exposure. Stress induction is typically performed on agar plates or in liquid culture (for review see [2]). Agar plates provide the benefit of eliciting “normal” behavior, with animals feeding regularly, moving in a stereotypical sinusoidal manner, and mating when partners are available [3]. However, these plate-based assays also suffer from two weaknesses. First, several environmental factors change over the duration of the experiment. Metabolic byproducts accumulate from the worms and bacteria, the biological state of the bacteria changes in response to the stressor which can have secondary effects on the worms, and the stressor itself can potentially degrade over the course of the experiment. While this effect can be minimized through periodic transfer of animals to new plates, physical manipulation introduces the possibility of additional stress or injury. Second, individuals frequently attempt to flee many of these environments and can be lost to “walling” on the sides of the petri plate, where they die from desiccation rather than from exposure to the stressor of interest. In contrast, liquid culture assays are performed with buffer-suspended animals in tubes or multi-well plates. For assays in tubes, sub-populations are usually sampled to determine survival status, while multi-well plates can be imaged similarly to agar plates [2]. However, animal behavior and physiology in liquid culture is quite different than on plates, with animals feeding less, altering egg-laying patterns, mating less, and engaging in bouts of vigorous swimming often described as “thrashing” due to its intensity compared to normal movement [3]. While liquid culture assays suffer from the same challenges of a changing environment that plate-based assays do, they do benefit from the elimination of sample loss due to walling.

Both traditional agar plate and manual liquid culture stress assays suffer from throughput and temporal resolution limitations imposed by the labor requirements of the assays. Additionally, most standard stress assays are performed on population cohorts without longitudinal information for individuals. To overcome these limitations, several technologies have been developed. For example, temporal resolution can be improved using automated scanner or camera based image acquisition and processing (e.g. *C. elegans* Lifespan Machine [4], WormScan [5], and worm tracking software [6]). Environmental standardization can be improved using microfluidic devices that allow constant profusion to flush out metabolic byproducts and replenish the stressor [7–10]. What is currently lacking, however, is a research platform that marries the benefits of automated image acquisition, distributed imaging, and the chemostat-like properties of flow through microfluidics with a device that allows longitudinal measurements of individuals in an environment that promotes “normal” plate-like behavior.

Here we present a new approach for studying stress response in *C. elegans* that combines the distributed imaging platform from the plate-based *C. elegans* Lifespan Machine [4], custom image processing and worm annotation software and a custom flow through microfluidic animal husbandry device. Our microfluidic device, the Stress-Chip, allows low population density (~1 animal per arena) animal culturing to facilitate high-throughput longitudinal studies. The single arenas use a structured artificial dirt [11–13] pillar arrangement to promote plate-like behaviors while maintaining the benefits of a flow-through aqueous environment. We find that the microfluidic platform can detect concentration dependent stress responses and can discern between genetic backgrounds. While developed for stress analysis, this research platform should be applicable to a variety of research questions, as survivorship is a fundamental measurement used in *C. elegans* research as diverse as anthelmintic-drug screening, aging studies, Alzheimer’s modeling, and toxicology studies.

## Results

### Design of the Stress-Chip Platform

To implement scanner-based microfluidic stress assays we designed a microfluidic chip (Fig 1 and S1 File) that addresses the physical constraints imposed by the scanner. In particular, the transparency unit (TPU) that provides transillumination casts shadows from the necessary fluid flow system that is attached to the chip and generates an exposure gradient perpendicular to the midline of the scanner. The exposure gradient is extreme enough that microfluidic devices centrally located on the scan bed are over exposed, rendering a narrow 5 mm to 90 mm “imaging zone” usable. By designing the arenas so that they are oriented in two parallel arrays (Fig 1A) we were able to fit a column of microfluidic chips in the imaging zone while maintaining a minimal scan width (~4 mm) in the orientation of the illumination gradient for each array scan. On this scale the gradient is unnoticeable and does not affect image thresholding and processing. The two inlets and two outlets are paired on each side of the design to consistently orient flow through the imaging zone to the middle of the scanner, minimizing the potential for shadowing due to the buffer inlet and waste outlet tubing (Fig 1).

**Fig 1.**
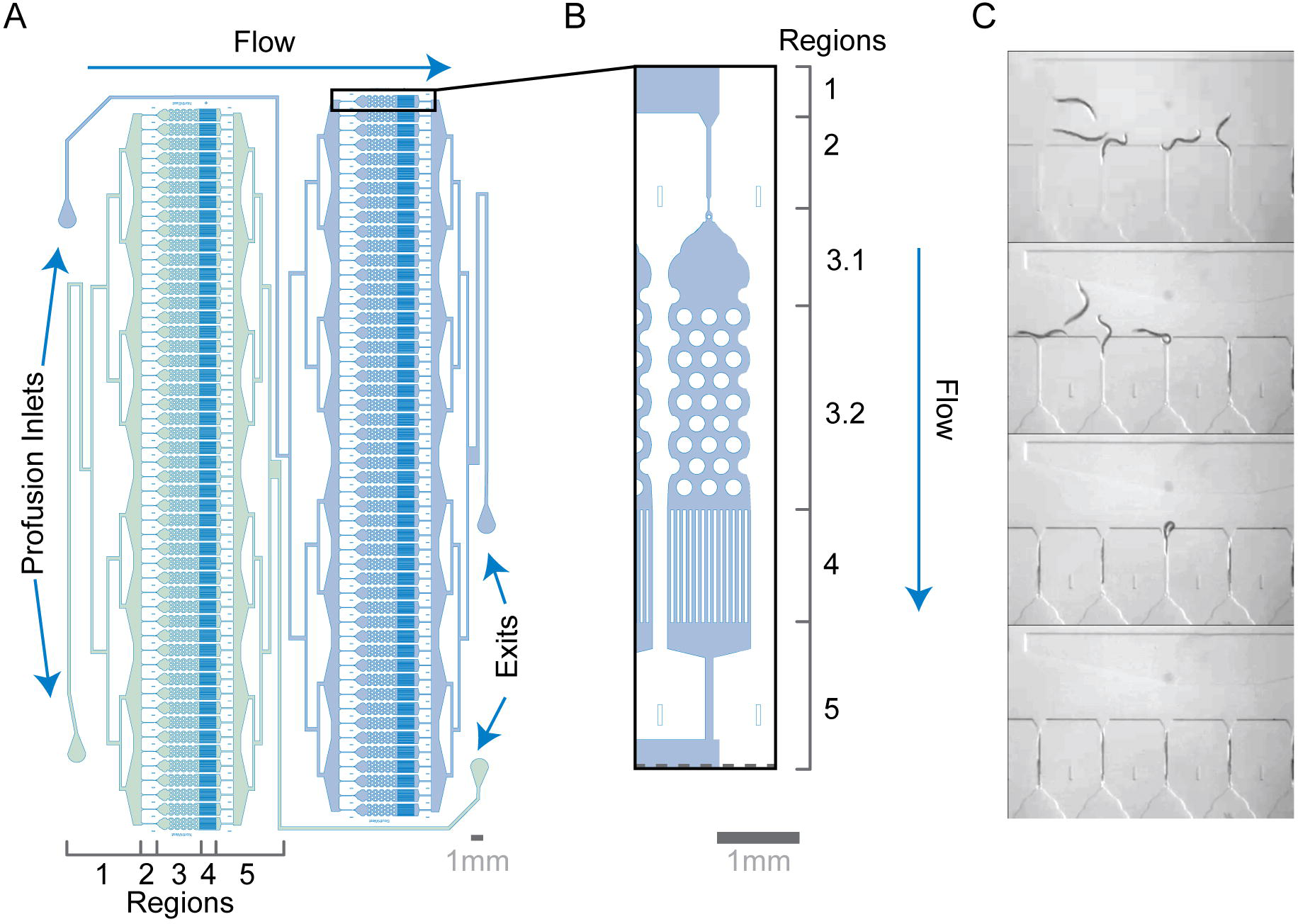
Microfluidic device for parallelized low-density worm culture. (A) The microfluidic chip is designed to have two arrays of 50 growth arenas. Flow through the system enters in region 1, a perfusion inlet on one side of the chip connected to a distribution network that ends in an open space upstream of the loading chambers. Flow then passes through the loading chambers (region 2) and enter the growth arenas (region 3). Flow exits each of the growth arenas through a filter (region 4) and reconnects to a single outflow network (region 5). The exit ports at the end of region 5 are located on the opposite side of the chip from the perfusion inlets to avoid shadowing by overhanging tubing. (B) Each arena includes a 2.2 mm x ~0.9 mm “crawl-zone” of 0.2 mm diameter pillars spaced 0.1 mm apart (see [12]) (region 3.2). Upstream of the pillar array is a “swim zone” (region 3.1) that stretches 1.1 mM from entry point to the first set of pillars. The combination of an entry plug and flow keeps loaded animals in the arenas. (C) Automated loading is facilitated by holding chambers with a 0.028 mm wide constriction points that enable the parallel pre-loading of 50 animals which are then moved into the chambers with a short burst of increased pressure (adapted from [10]).

In addition, the microfluidic device is designed to (1) reliably deliver worms to their respective arenas, (2) keep worms restricted to their specific arena for the duration of the experiment, and (3) elicit plate-like behavior while providing a constant profusion to deliver stressors and remove waste from the arenas. To accomplish this, we designed a device that can be divided into five regions with different functions. The first region consists of the profusion inlet and a series of bifurcations that form a distribution network that terminates at a large open space just upstream of 50 parallel loading channels (Fig 1A). The distribution network delivers worms to the loading channels during experimental setup and delivers buffer and stressor during the experiment. The open space enables self-distribution of the animals to the loading channels without requiring perfect distribution at the bifurcations of the distribution network (S1 Movie). The second region consists of an array of 50 loading channels that connect the open area at the end of the distribution network to the worm arenas. Each loading channel is a 0.05 mm wide channel sized to fit a single day-one adult *C. elegans* hermaphrodite that terminates in a 0.028 mm constriction point which precludes the animal from entering the arena without a burst of higher-pressure (Fig 1B and S1 Movie). This hold and burst loading approach has been successfully used previously [10]. The third region consists of an array of 50 arenas that house the animals for the duration of the experiment.

Each arena is 0.99 mm wide × 3.6 mm long at their widest points and feature two sub-regions. The first sub-region (swim-zone) is an open space that allows the worm to swim while preventing animals from crawling up into the distribution network by eliminating features that the worm can use to generate the force necessary to push through the loading channel constriction point (Fig 1B region 3.1). The second subregion (crawl-zone) is filled with a pattern of pillars that have been sized to generate plate like behavior in adult animals [11,12] (Fig 1B region 3.2). This combination of swim-zone and crawl-zone provides a more complex environment than previous microfluidic arrays for single worm husbandry [9,10,14]. Downstream of each arena is a series of 0.04 mm wide x 1.38 mm long channels spaced 0.035 mm apart that act as a filter that excludes adult worms from the exit, while allowing embryos to be flushed out of the chamber (Fig 1B region 4). The efficacy of loading and retention can be measured as the experimental yield, defined as the number of animals with either a call of death or identified as alive at the completion of an experiment relative to a max capacity of 50 arenas. For all experiments run to date using the system we have a yield of 68.8% (9,153 data points/13,300 arenas).

To provide a standardized environment, we attached the microfluidic chips to a pressure driven buffer system that can deliver the stressor to the animals and flush waste and progeny out of the arenas (Fig 2A). While the flow-through nature of the microfluidic chips helps to standardize the chemical environment, temperature standardization must also be accounted for. Nematodes are poikilotherms, and changes in temperature can have a profound effect on reproduction [15] and lifespan [16]. We therefore modified EPSON V800 flatbed scanners to reduce heat buildup from the internal components of the scanner. Previous scanner-based implementations used EPSON V700s, which use fluorescent bulbs instead of LEDs [4,5]. We found the V800s generate less heat, so we simplified the modifications that were made in the V700 *C. elegans* Lifespan Machine implementation [4]. In brief, the TPU lid remained unaltered, while the scanner base was ventilated in three locations (Fig 2B). The rear of the scanner base had a 125 mm x 25 mm opening, while both sides of the scanner had 75 mm x 50 mm holes cut approximately 75 mm from the back (Fig 2B). A small USB powered fan was then used to draw air from inside the scanner to maintain an appropriate surface temperature on the scan bed while avoiding direct airflow across the scanner bed (Fig 2B).

**Fig 2.**
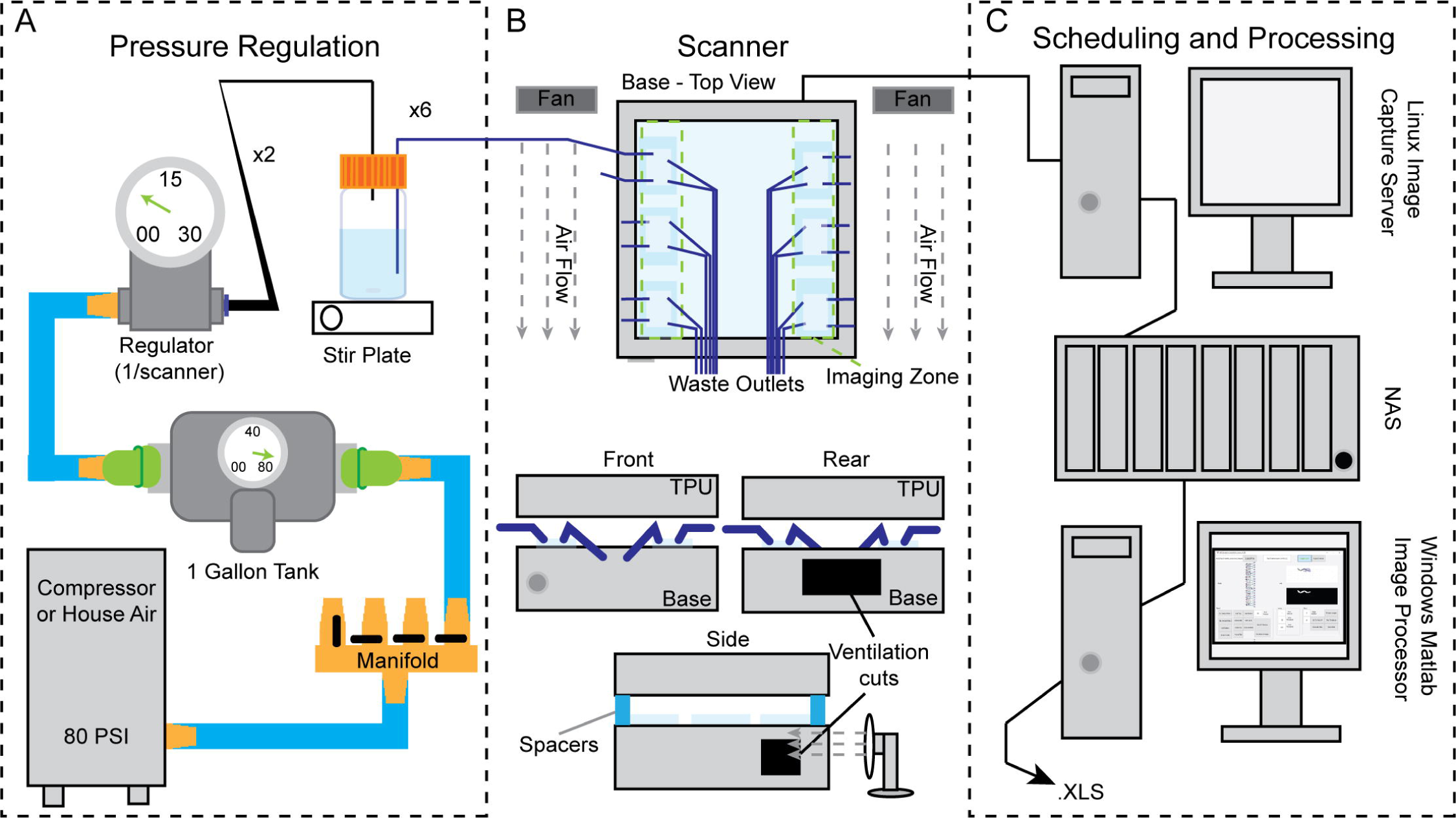
Scalable pressure drive and distributed imaging system. (A) Pressurized air was split through a manifold with ball valves to 1-gallon air tanks. Air from the 1-gallon tanks was regulated using a pressure-relieving micro-regulator to 1-5 psi. The regulated air was then passed through a t-split to two 1-liter bottles with custom fabricated lids that had seven 1.5 mm OD stainless steel stems that passed through the lid. One stem was used to connect the pressurized air to the air in the bottle, while the remaining six stems were connected internally to tubing that reached ~50 mm from the bottom of the bottle to allow for a stir bar, and externally to 1.5 mm ID tubing that ran to the profusion inlet ports of the microfluidic chips (see Fig 1A). (B) Microfluidic chips on 50 mm x 75 mm glass slides were mounted in the imaging zone approximately 12 mm from the edge of the scanner, with three chips on each side. (C) Image capture scheduling was performed using a Linux implementation of the *C. elegans* Lifespan Machine software [4] and images were stored on a local network attached storage device. Image processing was performed using custom MATLAB code (see materials and methods) run on a Windows 7 desktop.

To determine the temperature profile of the scanner bed during an experimental run, we developed a custom temperature recorder (S1 Fig and S2 File) with thermistor probes mounted on the same 50 mm x 75 mm glass slides that were used for manufacturing PDMS microfluidic chips. The temperature profile of the scanner bed was then determined by recording the temperature from 16 scanner bed locations every 1530 seconds during mock Stress-Chip experiments (Fig 3). We find that the experimental temperatures are consistent from run to run, and scanner to scanner (Fig 3C). Each 1-hour recording consists of a minimum of 120 temperature readings per probe, with each temperature reading representing the average of 50 samples taken in ~1 second (S2 file), from each of the 16 probes. While the average scanner temperature is constant across scanners and time, there can be temporal and spatial variation during an experimental run (Fig 3). The temporal pattern of the approximately 10-15-minute temperature cycle does not correlate with the 5-minute image sample collection, but rather tracks with the cycling of the HVAC unit that maintains temperature in the 20°C room in which experiments were performed. The mean temperature for each of the 16 locations also show differences, with a maximal departure from the scanner mean of ~0.5°C (Fig 3B).

**Fig 3.**
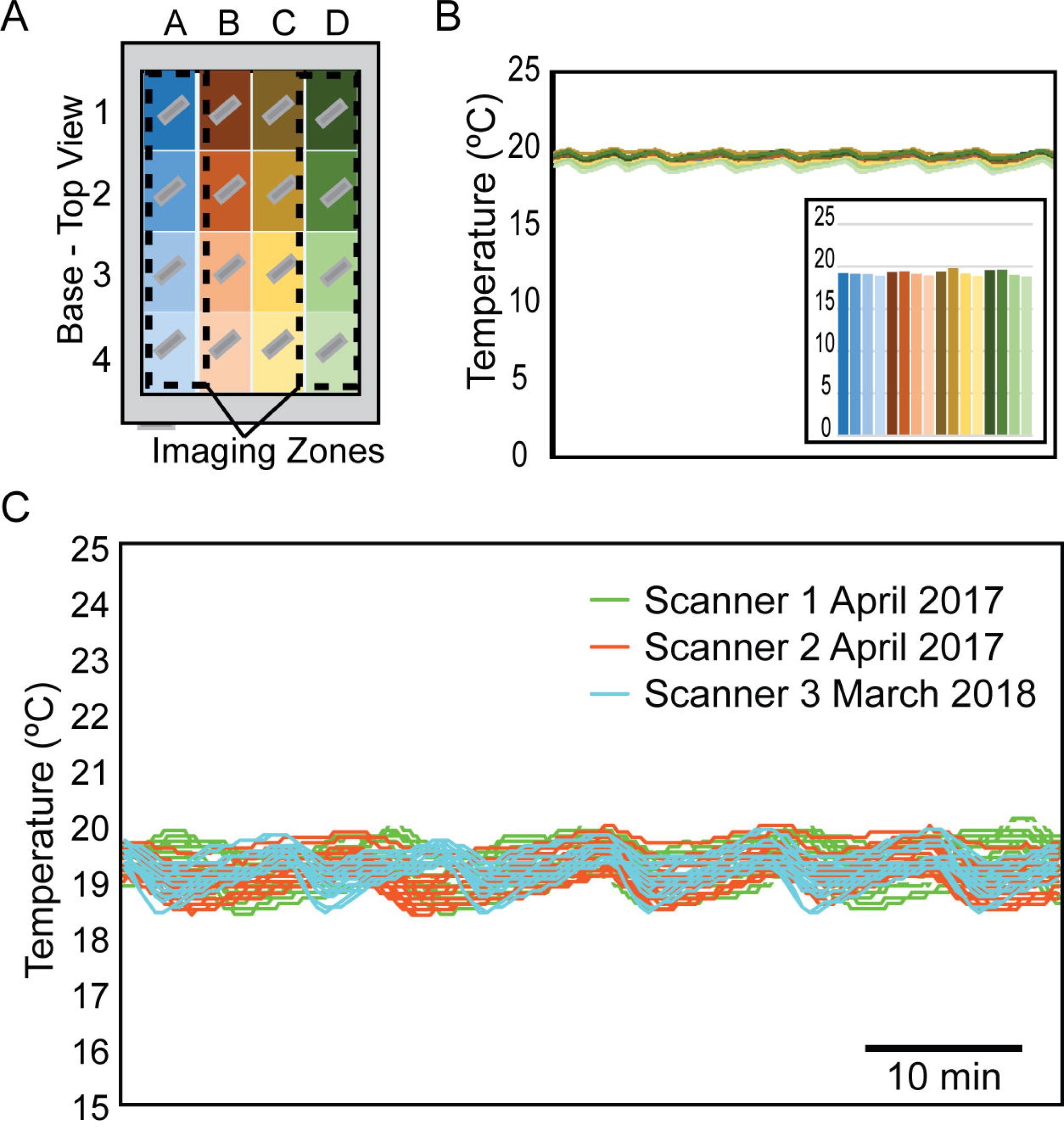
Scanner temperature stability. (A) Temperature probes were mounted on glass slides as PDMS chips are and arrayed to cover the entire surface of the scanner. The outermost columns of probes lie in the imaging zone where microfluidic chips would be mounted. (B) Temperature data from a mock run shows little variability in temperature over time. The cycling of temperature that is observed corresponds to HVAC cycling in the temperature-controlled room not with the imaging cycle of the scanner. (C) Measurements on different scanners on different days are consistent.

### Dietary Stress

The natural environment of *C. elegans* is decaying plant material [17,18], an environment marked by a boom/bust cycle of food that necessitates behavioral responses to address the absence of food, or low food densities relative to the worm population density [17,18]. One such response is the cessation of egg-laying in response to food scarcity during adulthood. The cessation of egg-laying results in matricidal internal hatching of progeny that subsequently use the mother as a food source. This response is known as facultative vivipary [19] or “bagging”. To determine if dietary stress would result in measurable matricidal bagging in a microfluidic environment we measured lifespan in M9, a salt buffer that lacks an energy source. The image quality from the scanners was adequate to identify matricidal bagging, with mothers losing opacity over time due to being internally consumed by their progeny (Fig 4A).

**Fig 4.**
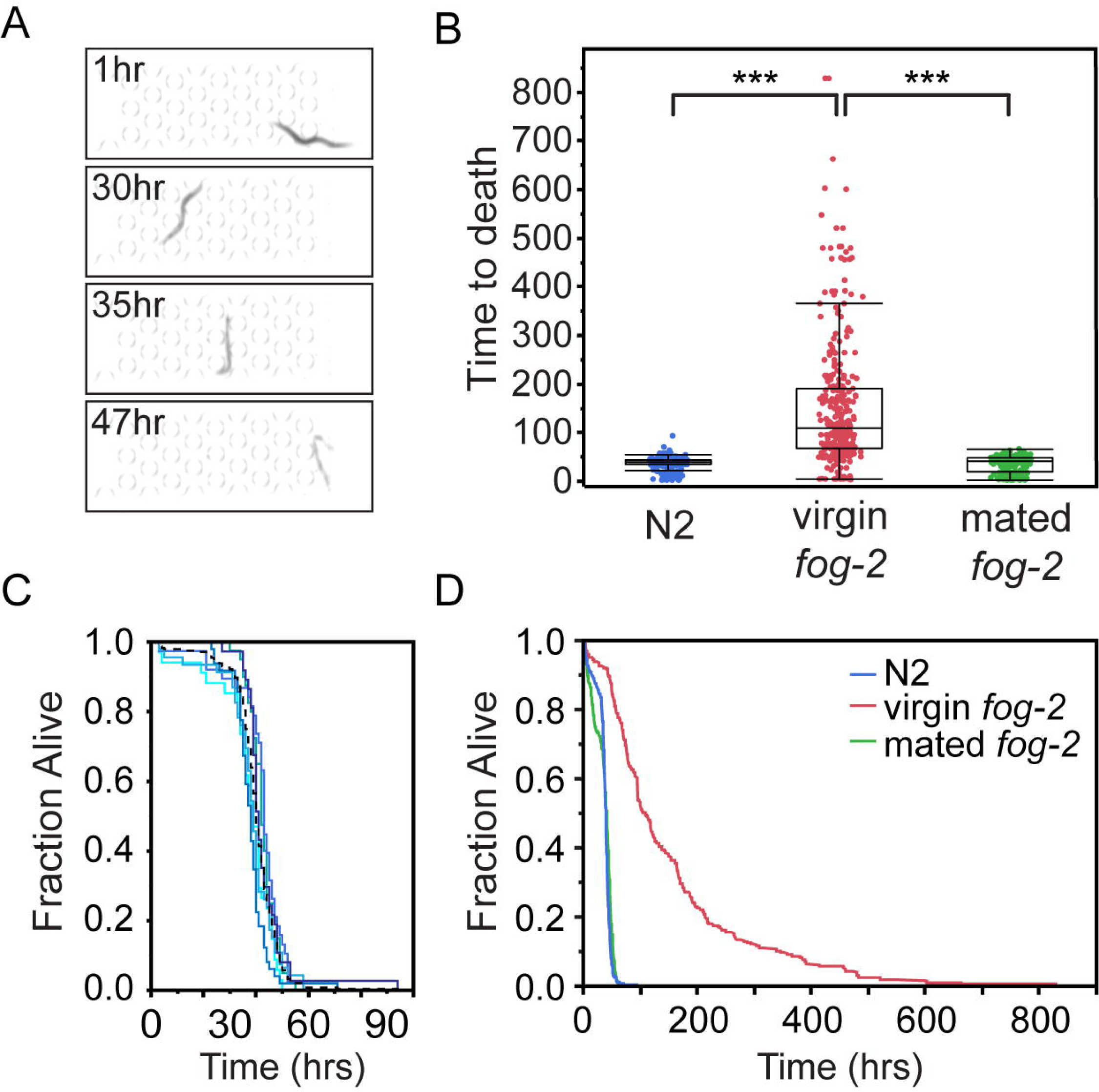
Dietary stress. (A) Images of N2 animal at 1, 30, 35, and 47hrs showing the characteristic “bag of worms” death. (B) Time to death for animals in M9 buffer (Number of animals = 244 (N2), 327 *(fog-2(q71))*, 263 (mated *fog-2(q71))).* *** signifies p<0.0001. The N2 versus mated *fog-2* comparison is not significantly different, with a p=0.87. (C) N2 Lifespan curves in M9 from day 1 of adulthood. Each solid line is a different sample lane representing between 34 and 49 individuals. The black dotted line represents the curve of all N2 animals. No significant differences are seen between replicates. (D) Lifespan curves in M9 from day 1 of adulthood. Log Rank p<0.0001.

Animals died quickly in the absence of food, with a median measured lifespan of 40 hrs (quartiles: 25%, 36.5 hrs; 75%, 44 hrs) and a mean lifespan of 39.9 hrs (SEM 0.6 hrs) (Fig 4B). The lifespan curves showed no significant differences between replicates, with curves generated for each lane showing similar behavior (Fig 4C). To further confirm that matricidal bagging was responsible for the observed deaths, we performed experiments in self-sterile virgin *fog-2(q71)* hermaphrodites that are unable to bag due to the absence of sperm [20]. Virgin L4 *fog-2(q71)* hermaphrodites were isolated and loaded into the arenas on the first day of adulthood. The sperm deficient animals greatly outlived the wildtype animals in the absence of food, with a mean lifespan of 152 hrs (Fig 4B), comparable to the lifespan of fed wildtype animals in other microfluidic devices (day 7 of adulthood for the life-span-on-a-chip worms [9] and day 7.3 for the WormFarm [7]). The prolonged lifespan is dependent on the absence of sperm, and mating *fog-2(q71)* hermaphrodites to males prior to loading in the Stress-Chips results in lifespans that are indistinguishable from wildtype (Fig 4B and D). To further validate our Stress-Chip platform we tested additional stresses in the absence of food using the *fog-2(q71)* background.

### Oxidative Stress

Oxidative damage naturally occurs as a result of metabolic processes, and it has been hypothesized that accumulation of oxidative damage is a fundamental driver of aging [21]. An organism’s environment can increase the rate of oxidative damage, either directly by the presence of oxidative species, or indirectly by increasing the rate at which internal oxidative species generation occurs. Understanding the relationship between the environment and an animal’s capacity to reduce oxidative species is fundamental to understanding aging and stress processes. *C. elegans* is a popular model for studying oxidative stress where oxidative stress can be introduced by hydrogen peroxide, paraquat, juglone, t-BOOH, arsenite, hyperbaric oxygen, and genetic mutation [2]. To determine the effectiveness of our platform for studying oxidative stress we generated lifespan curves for *fog-2(q71)* animals at 0.25 mM, 0.5 mM, 0.75 mM and 1 mM H_2_O_2_. The presence of H_2_O_2_ resulted in short lived animals that died in a manner qualitatively different than when death was induced by starvation alone. Animals in H_2_O_2_ showed a stereotypic rod-like death (Fig 5A) and retained opacity for the duration of the experiment, unlike worms that died due to matricidal bagging that lost shape and coloration. Our image processing software identified the timepoint at which movement ceased for each individual, and we generated dose dependent H_2_O_2_ lifespan curves of *fog-2(q71)* animals (Fig 5B). Interestingly, the curves generated from 0.5 mM and 0.75 mM showed variability in the effectiveness of the hydrogen peroxide. This suggests that the susceptibility curve for animals is steep in the 0.5 mM-0.75 mM range, and that minor variability in worm preparation and/or reagent preparation when assaying in that range can have a major effect on phenotype. To determine the LT50 response for hydrogen peroxide we determined the median lifespan of *fog-2* animals at the four concentrations of H_2_O_2_, revealing a negative exponential relationship between lifespan and H_2_O_2_ concentration (Fig 5C).

**Fig 5.**
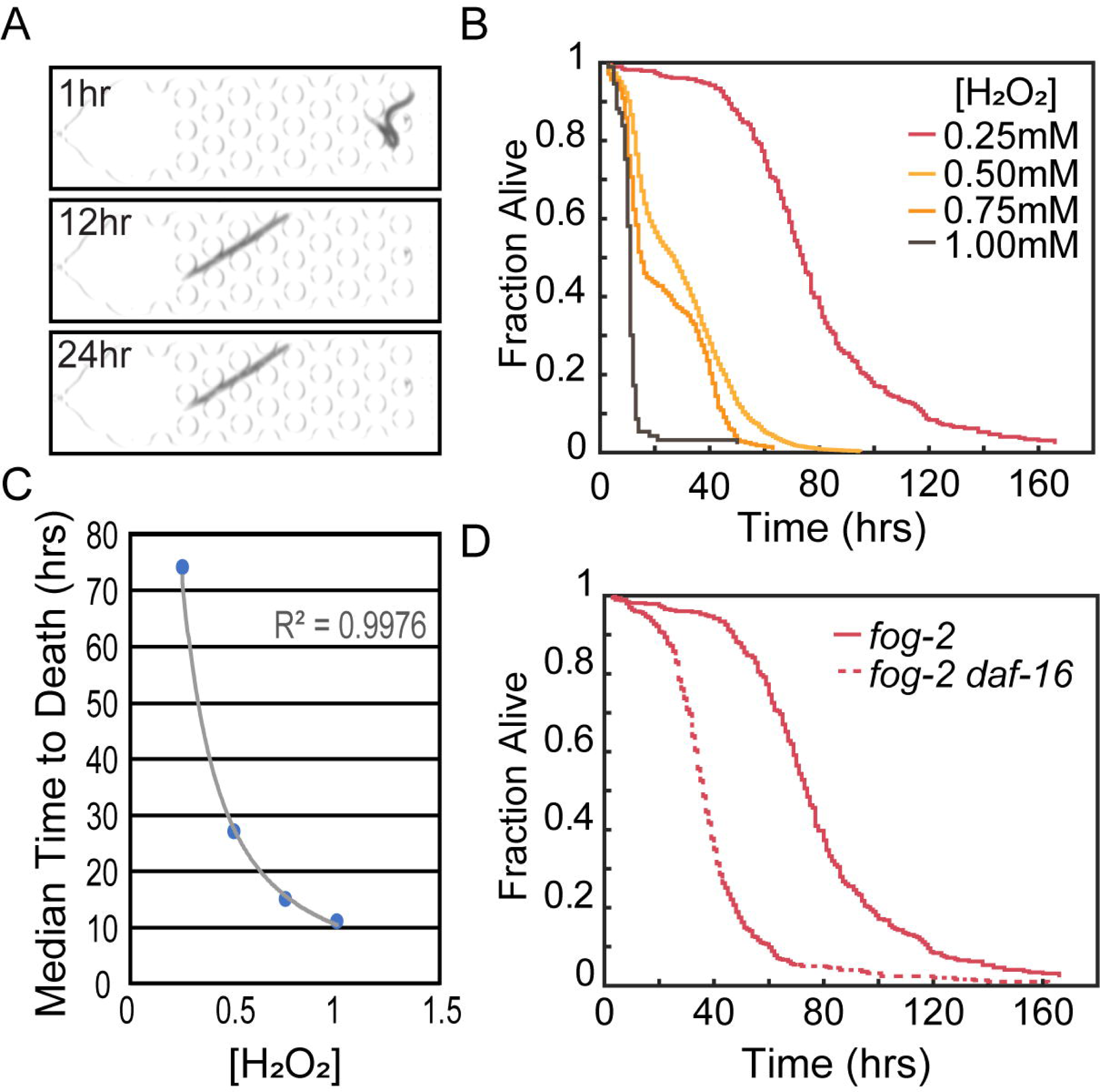
Oxidative Stress. (A) Images of a representative *fog-2(q71)* animal at 1, 12, and 24hrs showing the characteristic rod-like death (B) Lifespan curves for *fog-2(q71)* animals in M9 buffer supplemented with increasing concentrations of H2O2 (Number of animals = 379 (0.25 mM), 3518 (0.5 mM), 276 (0.75 mM), 93 (1 mM)). (C) Power fit of median lifespan versus concentration of H_2_O_2_ yields an equation (y = 10.503x^−1394^) with an R^2^ of 0.9976. (D) Comparison of *fog-2(q71)* and *fog-2(q71) daf-16(mgDf47)* animals at 0.25 mM H_2_O_2_ (Number of *fog-2 daf-16(mgDf47)* animals is 498). Log-Rank *p<0.0001.*

We next determined if the Stress-Chip platform could detect hyper-sensitivity to hydrogen peroxide. Previous work has shown that the insulin signaling pathway is required for wildtype oxidative stress response [22,23]. The primary transcriptional effector for insulin pathway dependent oxidative stress response is the transcription factor DAF-16, which responds to oxidative stress by translocation to the nucleus where it can activate stress response genes [24]. H_2_O_2_ in particular has been shown to drive DAF-16 to the cytoplasm in an *age-1* dependent process [25]. We therefore generated lifespan data for *fog-2(q71) daf-16(mgDf47)* animals in the presence of 0.25 mM H_2_O_2_. Loss of *daf-16* resulted in faster death (Fig 5D), with a median lifespan of 36 hours compared to 74 hours for control animals. When compared to the sensitivity of *fog-2(q71)* animals, loss of *daf-16* results in animals which are approximately 65% more sensitive to H2O2, with *fog-2(q71) daf-16(mgDf47)* responding to 0.25 mM H2O2 as *fog-2(q71)* control animals would be expected to respond to 0.413 mM H_2_O_2_.

### Osmotic Stress

Hypertonic conditions can drive protein aggregation [26], dsDNA breaks [27], and accelerated aging [28]. We therefore tested the efficacy of the Stress-Chip at measuring *C. elegans* lifespan under varying sodium chloride concentrations. Under standard laboratory conditions worms are grown on NGM which has 21 mM NaCl. To present hypertonic conditions we used M9 buffer (typically 86 mM NaCl) adjusted to 300, 400, and 500 mM NaCl because wildtype worms show a robust ability to survive NaCl concentrations ≤200 mM [29,30]. Worms in high NaCl conditions exhibit a qualitatively different phenotype than those under starvation or H_2_O_2_ conditions. As previously observed, animals begin with a squat phenotype at the time of the application of high salt, and regain length over the course of the experiment (Fig 6A) [30]. This response is presumed to be mediated by upregulation of glycerol biosynthesis [30,31]. Despite these changes in length, our longevity-detection software remains robust at determining the time at which movement ceases, allowing us to generate lifespan curves of *fog-2(q71)* animals under varying concentrations of NaCl.

**Fig 6.**
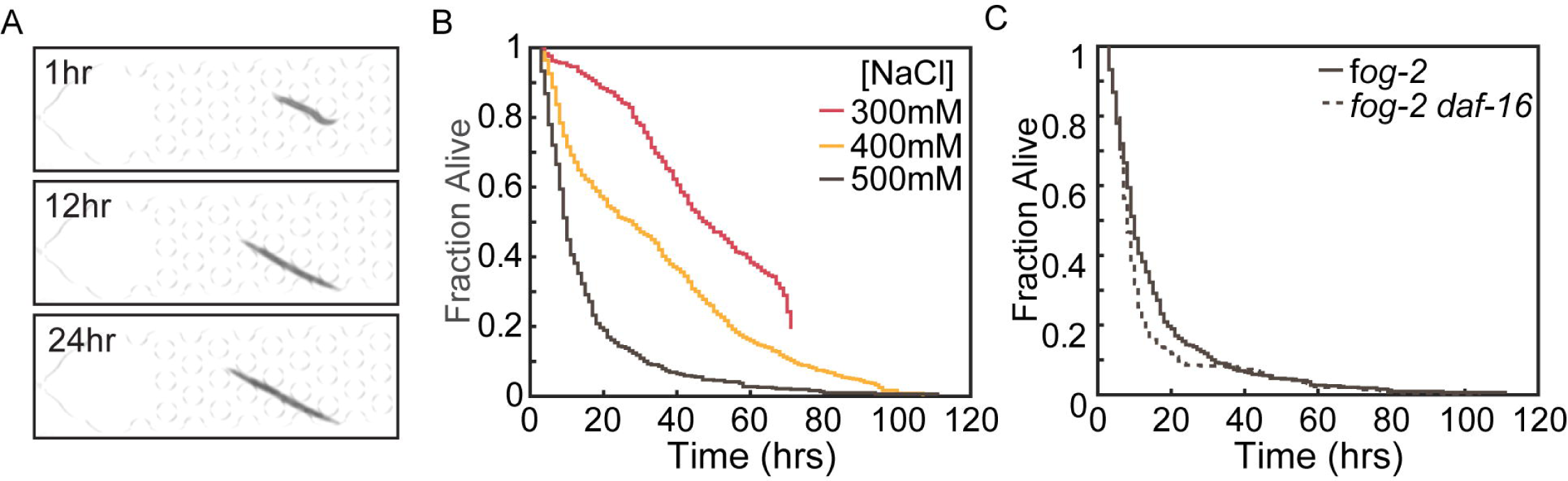
Osmotic Stress. (A) Images of a representative *fog-2(q71)* animal at 1, and 24hrs showing the characteristic squat worm phenotype at NaCl application, and recovery over time. (B) Lifespan curves for *fog-2(q71)* animals in M9 buffer supplemented with increasing concentrations of NaCl (Number of animals = 297 (300 mM), 792 (400 mM), 508 (500 mM)). Log-Rank *p<0.0001.* (C) Comparison of *fog-2(q71)* and *fog-2(q71) daf-16(mgDf47)* animals at 500 mM NaCl (Number of *fog-2(q71) daf-16(mgDf47)* animals is 450). Log-Rank *p=0.0005.*

Animals showed a concentration dependent decrease in lifespan in response to NaCl (Fig 6B). Our results are comparable to those obtained by others in plate-based assays. For example, we observe a mean lifespan in 400 mM NaCl of 33.2+/− 0.75 hours for day 1 adult animals. Others, using day 4 adults in the presence of food, observe a mean lifespan in 400 mM NaCl of 40.3+/− 6 hours [32]. Previously published work with day 1 adult animals showed a wide range of lethality 24 hrs post exposure to 400mM NaCl, from >90% lethality [29] to <20% lethality [26]. Our observation of 48.6% lethality at 24 hours for 400mM NaCl lies in the range. In previously published work using a single time point, 500mM NaCl resulted in ~100% lethality at 24 hours [33], or 97% [31] at 24 hours. Our microfluidic approach gives similar results at 500 mM NaCl, with a mean lifespan is 15.4 +/− 0.75 hours, and 85.1% lethality at 24 hours (Fig 6B). These results support the notion that microfluidics-based stress assays using *fog-2* animals in the absence of food give results comparable to those of wildtype animals on agar plates seeded with bacteria.

To compare our NaCl results with our H_2_O_2_ results, we tested *fog-2(q71) daf- 16(mgDf47)* animals in 500 mM NaCl. Previous work has shown that in an otherwise wildtype background, *daf-16* is not required for acute or adaptive osmotic stress response when animals are grown on high NaCl plates [29]. This is in contrast to *age-1* or *daf-2* animals, which are resistant to elevated NaCl concentrations in a *daf-16* dependent manner [29]. The resultant *fog-2(q71) daf-16(mgDf47)* lifespan curves showed a negligible difference in lifespan when compared to *fog-2(q71)* controls (Fig 6C). These results support the view that osmotic regulation is different from other stress pathways that are regulated in wildtype animals by a limited number of master regulators (e.g., DAF-16 and SKN-1) [34,35]. Additionally, these results demonstrate that the Stress-Chip platform can detect genetic sensitivity in a stress specific manner.

## Discussion

Here we present validation of the Stress-Chip, a research platform that is a novel fusion of microfluidic and scanner-based approaches with potential use in a broad range of research applications. We find that the Stress-Chip is applicable to the study of a variety of stresses, including dietary, osmotic, and oxidative stress. The system can effectively establish dosage responses, distinguish sensitivity between genetic backgrounds, and show stress-specific enhancement of susceptibility for genetic backgrounds; all key components in large-scale *C. elegans* stress studies. The success of the Stress-Chip is built on five key features: (1) A high temporal resolution for determining time of death, (2) chemostat-like properties of a flow through system, (3) a complex microfluidic environment that can promote plate-like behaviors, (4) temporal control of stressor application, and (5) easy scaling. In addition to those benefits, the system enables testing of a variety of stresses while minimizing experimenter time and effort.

To achieve a high temporal resolution, the Stress-Chip capitalizes on the distributed imaging platform of the *C. elegans* Lifespan Machine. The scanner and image acquisition software provide a one hour temporal resolution while collecting images continuously throughout the entire lifespan of the animals [4]. This provides an opportunity for greater informational depth than usual single-timepoint stress assays. While single time points (e.g. 24-hour survival) can be used to generate LD50 information for compounds [2], continual monitoring also allows LT50 measurements and lifespan curve analyses to be performed [16] (e.g. Fig 5C).

In addition to providing a high temporal resolution, the scanner-based approach of the Stress-Chip platform is also easy to scale after initial setup due to the overcapacity of most pressurized air and computer systems. The low volume of compressed air used (less than 1-liter at 5 psi per scanner-day) is orders of magnitude below the capacity of typical air compressors or house air systems, and a typical modern Linux computer system can run many scanners. For example, our lab currently runs 20-25 scanners for Stress-Chips and plate-based assays on each Linux computer running the *C. elegans* Lifespan Machine software. The platform therefore allows easy addition of capacity after the initial set up of pressurized air and computer systems, by acquisition of additional scanners. Each scanner represents a potential throughput of 600 animals, while one computer system and pressurized air system can potentially handle 20+ scanners for a potential throughput of 12,000 animals.

Previous scanner-based stress-assays have been performed using plates or wells to house animals [4,5]. Typical scanner-based assays require the plates to remain stationary for the duration of the experiment [4], which eliminates the possibility of moving animals to new media to avoid changing environmental conditions. Microfluidic animal husbandry provides a solution to the problem of environmental change by maintaining a chemostat-like environment while allowing fine control over the delivery of stressors. This benefit is not unique to the Stress-Chip, and is a feature of other nematode-housing microfluidic devices like the WormSpa [14], Lifespan-on-a-Chip [9], and WormFarm [7]. Flow through microfluidic animal husbandry has additional benefits including the opportunity for size-dependent flushing of progeny from the arenas. With this arrangement, wildtype animals can be cultured without chemical sterilization by compounds like the anti-cancer drug 5-Fluoro-2′-deoxyuridine (FUdR). FUdR exposure is a frequently used method when studying adult hermaphrodites [36], but it is known to change lifespan and stress resistance (e.g. osmotic stress [37]). While all flow-through microfluidic devices offer chemostat-like benefits, our device differs from other platforms by being the first optimized for scanner-based imaging. Additionally, our device houses worms individually (versus populations for the WormFarm [7]) in arenas that can be run for weeks (versus 24 hours or less for the WormSpa [14]) while being easy to load and a more complex spatial environment (versus simple arenas with no autoloading for the Lifespan-on-a-Chip [9]). Additionally, the arenas in our device eliminate stress-induced escape attempts that result in walling in plate-based assays.

The high temporal resolution, chemostat-like properties, environmental control of escape behavior, ease of stressor application, and ready scalability of the Stress-Chip make it a highly adaptable research tool. The ease of controlling flow-through microfluidics also allows fine temporal control of the environment, with stream switching enabling a variety of research paradigms like controlled duration exposure, pulse-chase, or cyclic stressor application that would be difficult to achieve in plate-based assays. Additionally, we believe that the Stress-Chip platform holds promise for many research areas beyond stress response. The general applicability of time-to-death measurements makes it suitable for anthelmintic-drug screening, aging studies, Alzheimer’s modeling, and toxicology studies.

## Materials and Methods

### Reagents

All experiments were performed in M9 Buffer (22 mM KH_2_PO_4_, 42 mM Na_2_HPO_4_, 86 mM NaCl, 1 mM MgSO_4_) sterilized by autoclaving [38], or M9 Buffer supplemented after autoclaving with H_2_O_2_ (Fisher H325) or NaCl (Fisher BP358212) to achieve the noted concentration. For profusion through the microfluidic devices all solutions contained 0.01% triton (Sigma T8787) and were continually mixed at low speed using a stir plate/stir bar for the duration of each experimental run.

### Animal Husbandry and Worm Preparation

All animals were maintained at 20°C on NGM plates seeded with *Escherichia coli* OP50 [39]. *C. elegans* strains were obtained from the Caenorhabditis Genetics Center (CGC). Strains used in this study were N2 [39], CB4108 *fog-2(q71)V* [20], and AU166 *daf- 16(mgDf47) I; fog-2(q71) V* [40]. Synchronized worm populations were obtained by hatching sodium hypochlorite axenized embryos in M9 buffer [38]. Hatched worms arrested as L1s until they were transferred at a concentration of 2000 worms per plate onto 100 mm NGM plates seeded with OP50. All populations were maintained at 20°C throughout development. For experiments with virgin *fog-2(q71)* feminized hermaphrodites, *fog-2(q71)* hermaphrodites were separated from males prior to adulthood. This was achieved by rinsing the age-synchronized populations from the NGM plates two days post L1 plating. Worms rinsed from the plate with M9 were collected with a glass pipet and centrifuged in a 15 mL conical at 3200 rpm for 2 min. All but approximately 1 mL of supernant was removed. The conical was then topped off with M9 and centrifuged again to rinse away any residual bacteria. Approximately 4 mL of the resultant worm suspension was moved to a glass petri dish. Capillary tubing was heated and pulled to form a fine needle. The needle point was then broken to create an opening large enough to capture an L4 worm. The capillary was then mounted in a gasket with flexible tubing and operated by mouth pipetting. Light suction was applied to select unmated hermaphrodites from the worm suspension. This form of selection is more gentle [41], faster, and more efficient than pick selection. Once a large enough population of L4s was collected, they were spun down briefly at 3200 rpm and plated onto a fresh NGM plate seeded with OP50. The plate was then placed into the incubator to allow the worms to spread out. After the worms had sufficiently separated, the plates were searched, and rare carry-over males were removed. The plates were checked for males three times to ensure that no mating could occur. Plates were then incubated at 20°C to allow the L4 animals to reach adulthood.

Selected day 1 adults were rinsed off NGM plates with M9 and collected by glass pipet. Worms were then centrifuged in a 15 mL conical at 3200 rpm for 2 min. All but approximately 1 mL of supernant was removed. The conical tube was then topped off with M9 and centrifuged again to rinse away any residual bacteria. The supernatant was again removed until approximately 1 mL remained. Worm concentration was determined worm counts from 10 μL samplings. Each array held 50 worms and there were 12 arrays per scanner run requiring a total of 600 worms. To facilitate loading, ~60 worms were aliquoted into each of twelve 1.5 mL microcentrifuge tubes. To load the microfluidic chips the worms were pipetted from individual microcentrifuge tubes onto the loading port of a chip. They were then encouraged to move from the surface of the chip into the loading port by hand manipulation with a loop tool. Once in position a syringe filled with M9 + 0.01% triton was connected by polyethylene tubing to the loading port with a 1.5 mm stainless steel tube. Soft hand pressure was applied to distribute the worms across the array of loading chambers and then pressure was increased to force the animals into the arenas. Backflow was then applied to clear any worms that were not properly loaded into the arenas. This was repeated twice per chip. Once a chip was completed it was placed into position on the scanner.

### Microfluidic Device Fabrication

Microfluidic chips were created using standard soft lithography techniques [42,43]. In brief, chip designs were drafted in Vectorworks Fundamentals (Vectorworks, Inc.) and photomask transparencies were printed at 20k resolution (CAD/Art Services Inc). CAD files are available for download at https://blogs.uoregon.edu/phillipslabmicrofluidics/. The mask was then translated into a master using SU-8 (MicroChem Corp.) photolithography. Microfluidic devices were then made using Polydimethylsiloxane (PDMS) (Sylgard 184, Dow Corning Corporation) at the standard weight ratio of 10 elastomer:1 curing agent. 1.5 mm holes were punched using biopsy punches to gain access to the patterned channels, and the PDMS devices were bonded to 50×75 mm glass slides by air plasma exposure (PDC-32G Plasma Cleaner, Harrick Plasma Inc.).

### Image Acquisition

Images were acquired using the *C. elegans* Lifespan Machine software [4] previously developed for petri plate based assays. Microfluidic devices were scanned in grayscale at 3200 dpi and a color bit depth of 16, and the resultant images were stored as Tiff files for further processing. Scans of the 50-arena arrays were initiated every 5 minutes to achieve a scan frequency of 1-scan/hour for each of the 12 arrays. Arena location and time of scan were stored in the file metadata.

### Image Processing

Image processing was performed using custom MATLAB code (see S4 File). Image stacks were loaded and aligned according to user selected reference points using a custom MATLAB GUI (S2 Fig and S3 File). These reference points were then used to segment the array into the 50 individual arenas. Worm detection was then performed through a two-step process. First, each individual arena was cropped from the larger image and then processed using MATLAB’s image processing toolbox. The cropped arenas were converted to black and white images and then all objects too small to be worms were removed (300-pixel threshold by default, but user adjustable). Then, the size, location and dimensions of each potential worm-like object were calculated. Second, starting at the end of the experiment at which point all animals should be deceased and unmoving, and working to the beginning, the code cycles through the timeline of each individual arena. Using the object properties previously calculated, the relative locations and sizes of object centroids in adjacent image frames for each arena are compared. The determination of live/dead was made by determining if the worm object was seen to shift more than our threshold (7 pixels) between two images. This was repeated for each worm object in the arena at time of processing. However, because it was possible for the device to be disturbed and a global frameshift to occur between two images, an additional round of processing was performed in which the presence or absence of movement was tracked over successive frames. This allows the user to threshold the alive/dead call based on multiple observed movements rather than a single observation. Afterwards, a text file with the number of moving worms in each arena at each timepoint and a MATLAB workspace with all the metadata (location, worm objects, etc.) for each arena is saved. Further annotation can be performed during data analysis and any outliers observed can be manually checked through reloading the metadata data file and observing the time lapse of the outlier’s arena.

### Statistical Analysis

The raw MATLAB output file was reformatted, and the worm counts for each arena were translated into death times. To avoid spurious death calls from random movement late in the experiment, time of death was defined as the last time point before 4 sequential hours with no observed movement. Animals that remained alive at the completion of the experiment were censored at the time the experiment ended and were included in the analysis as such. The resultant lifespan curves were analyzed using JMP Statistical Software (SAS Institute Inc.). Lifespan curves presented are Kaplan-Meier curves, and statistical differences were determined by log rank test. Mean lifespans were compared by ANOVA analysis with posthoc Tukey-HSD.

## Acknowledgements

*C. elegans* strains were provided by the CGC, which is funded by NIH Office of Research Infrastructure Programs (P40 OD010440). We would like to thank the Phillips lab members for helpful discussion, Daniel Steck for technical advice on developing the temperature recorder, the Lu Lab at Georgia Tech for technical advice on pressure driven microfluidics, and the University of Oregon’s Lokey Laboratory nanofabrication facility for equipment and technical assistance.

## Supporting information captions

**S1 Fig. Parallel Temperature Recorder.** (A) The temperature recorder was built using a Raspberry Pi 3 single board computer, 16 thermistor probes, a custom printed circuit board and a custom case that provides separation between the analog and digital components for reduced electrical interference. (B) The custom PCB holds two 8 channel MCP3008 ADC chips. Signal noise was also reduced by inclusion of bypass capacitors C1-C3 and C4-C6. (C) A custom MATLAB GUI enables automated temperature experiments to be run at several discrete sample rates ranging from 2 samples/min to 4 samples/hour.

**S2 Fig. Semi-Automated Image Processing Interface.** (A) Global image alignment pane. Using user set reference points, the interface corrects for any small deviations in the relative angle between chip and scanner. (B) Arena segmentation occurs after the user selects the bounds of the arena as well as number of arenas in each viewing frame. This also allows the user to exclude parts of the device from analysis that may have become clogged during operation as well as makes the interface adaptable between different variations of arena design. (C) Worm size and data range thresholding is set by the user and allows the exclusion of timepoints where image quality might have been lost through clogging. It also allows for the size threshold to be altered in case different stages or strains of worms necessitate changes in the image processing algorithm. (D) Individual worm viewing, and playback allows the user to check any single chamber at any timepoint in case there are any concerns about the worm inside or any discrepancies that require investigation. (E) The data review and export panel allows the user to view timelapses of the data and displays both the unprocessed image as well as the computer vision output. In addition, once the user is satisfied with thresholding parameters, this panel is also used to direct the interface to generate and export the final data file with annotated worm deaths.

**File. Printable Stress-Chip Design.**

**File. MATLAB Temperature Recorder Software and Users Guide.**

**File. MATLAB Image Processing Code.**

**File. Stress-Chip Data.**

**S1 Movie. Autoloading of the Stress-Chip.**

